# The extra-embryonic space is a geometric constraint regulating cell arrangement in nematodes

**DOI:** 10.1101/2021.06.12.448179

**Authors:** Sungrim Seirin-Lee, Akatsuki Kimura

## Abstract

In multicellular systems, cells communicate with adjacent cells to decide their positions and fates. Cellular arrangement in space is thus important for development. Orientation of cell division, cell-cell interaction (i.e., attraction and repulsion), and geometrical constraints are the three major factors that define cell arrangement. Here we found that the amount and location of extra-embryonic space (ES), the empty space within the eggshell not occupied by embryonic cells, are critical to define cell arrangement in the 4-cell stage embryo of nematodes. This discovery was motivated by observations of a T-reversed-type arrangement, which was not explained by a model assuming simplified shapes of the eggshell, in our previous experiments. In this study, we incorporated the precise shape of the *C. elegans* eggshell in our newly developed multicellular morphology model based on the phase-field method. The new model succeeded in reproducing the T-reverse arrangement, demonstrating the importance of the precise shape of the eggshell. Further analyses revealed that the amount and location of ES is critical to develop various cell arrangements. Overall, our analyses characterized the roles of new geometrical contributors to cell arrangements, which should be considered for any multicellular system.

## 1 Introduction

Arrangement of cells which defines how cells contact each other is important in developmental processes. It mediates correct cell-to-cell communication and ultimately determines specific cell fates and body plan (Gilbert and Michael 2019). This is one of the homeostasis mechanisms in multicellular organisms including humans (Shahbazi 2020). The mechanisms determining cell arrangement can be classified into three factors: orientation of cell division, interaction (repulsion and attraction) between cells, and geometrical constraints provided by surrounding structures where cells are confined such as the eggshell (Baena-López et al. 2005; Gloerich et al. 2016; Yamamoto and Kimura 2017). Numerical models including these factors have successfully explained cell arrangements during the early embryogenesis of *C. elegans* and sea urchins (Kajita and Yamamura 2002; Akiyama et al. 2010; Schulze and Schierenberg 2011; Fickentscher et al. 2013; Pierre et al. 2016; Yamamoto and Kimura 2017).

Nematodes are good biological models to study cell arrangement during embryogenesis (Dolinski et al. 2001). In particular, the 4-cell stage of nematode embryos shows simple and diverse types of cell arrangement, which makes this stage a good target to comprehensively understand the mechanisms underlying cell arrangement. The P0 cell refers to the 1-cell stage after fertilization in *Caenorhabditis elegans*; it divides asymmetrically into two different daughter cells, the P1 cell and AB cell (2-cell stage). The AB cell first divides into ABa and ABp cells, and the P1 cell then divides into EMS and P2 cells (Gönczy 2005). As the P2 cell is adjacent to the ABp cell, but not to the ABa cell, the fate of ABp is distinct from that of ABa because of the signal from the P2 cell (Bowerman et al. 1992; Mickey et al. 1996). In our previous study (Yamamoto and Kimura 2017), we constructed a model for cell arrangement at the 4-cell stage considering three factors (cell division orientation, repulsion and attraction between cells, and the ellipsoidal eggshell as a geometrical constraint). This model succeeded in reproducing the four types of cell arrangement observed in different species, and in *C*. elegans individuals with different aspect ratios (ARs) of the ellipsoidal eggshell. The four types of cell arrangements were named as Pyramid, Diamond, T-shaped, and Linear. The study showed that the asymmetric attraction between cells plays an important role in improving the robustness of cell arrangement, whereas the eggshell AR is a source of diversity in cell arrangement. The model that we constructed preciously was named as the asymmetric attraction (AA) model, based on vertex mechanistic dynamics between the mass points of cells. The study also found another type of cell arrangement when the attraction between cells were impaired, which we named as the T-reverse type (Fig. 1A). The T-reverse arrangement could not be explained by the AA model, suggesting an important missing component in the AA model, and thus in our current understanding of cell arrangement.

**Figure 1:**
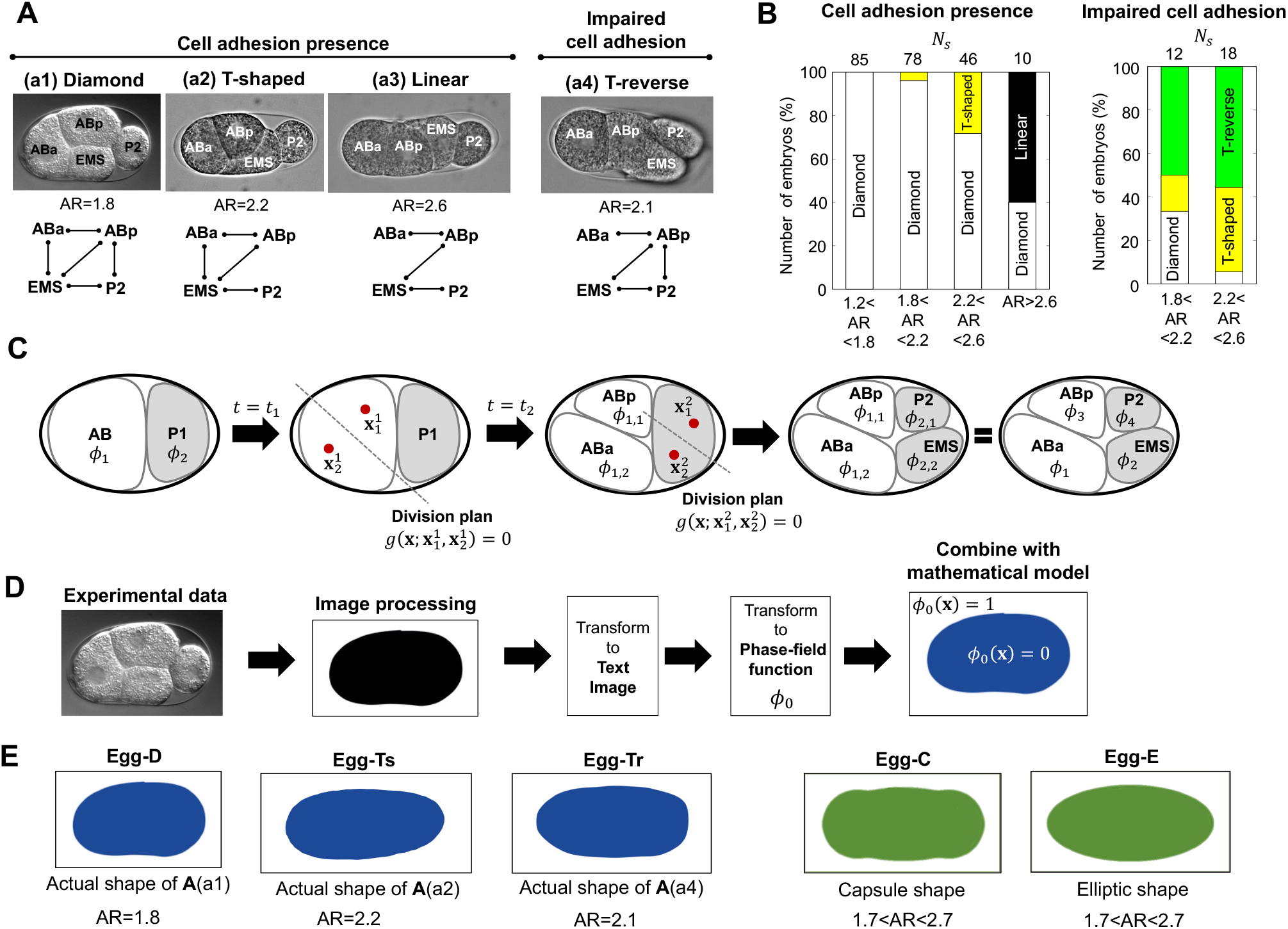
Types of cell arrangement and mathematical model by using the phase-field method. (A) Cell arrangement in the 4-cell stage of the *C. elegans* embryo as observed in experiments. The lower panels of the cell networks show a cell-to-cell contact state. AR indicates the aspect ratio. ‘Impaired cell adhesion’ indicates the embryo with knockdown of *hmr-1* and *hmp-2* genes, which are involved in cell adhesion. The normal (wild-type) embryo has an AR of ∼ 1.7, and embryos with a larger AR were obtained from the mutation of the *lon-1* gene and/or knockdown of the *C27D9*.*1* gene. (B) The types of cell arrangement that appeared in each AR range. *N*_*s*_ indicates the sample number of embryos. (C) Schematic diagram of modeling. The red dots indicate the spindles and the dotted line is the division plane determined by the location of spindles. (D) The process of converting the actual eggshell shape to a phase-field function. (E) The eggshell shapes that we used in the numerical experiments. Egg-D, Egg-Ts, and Egg-Tr were generated using the process (D) with the actual eggshell experimental data from (A). Egg-C and Egg-E were artificially generated using a phase-field model (Materials and Methods Eq. (2a)).

To determine the missing component, we focused on the third major factor influencing cell arrangement, namely, geometrical constraints. The effect of geometrical constraints has been less explored in the study of cell arrangement. For example, the eggshell has been simplified in mathematical models. Indeed, the AA model was based on a simple ellipsoidal eggshell. Therefore, we hypothesized that the limitation of the AA model is attributed to the simplification of the eggshell. To prove our hypothesis, in this study, we constructed a multicellular morphology model by using a multi-phase-field modeling method by which we can describe precise eggshell shapes. We then explored the cell arrangements at the 4-cell stage of nematodes: Diamond, Tshaped, T-reversed, and Linear types (Fig. 1A). In particular, we incorporated the real eggshell shape by combining imaging data with phase-filled modeling. Using this multicellular morphology model, we succeeded in reproducing the T-reverse arrangement and found that the precise shape of the eggshell affects the cell arrangement.

From the investigation of the reason for such sensitivity, we found out that the ‘empty’ space within the eggshell, which is not occupied by the cells but is filled with extra-embryonic matrix, can play a critical role in determining the cell arrangement. We named the empty space as the extra-embryonic space (ES). We further demonstrated that the variability in the amount and location of the ES can induce variability in cell arrangements even with the same condition of cell division orientation, cell-cell interaction, and the AR of the eggshell. Moreover, we revealed that the effect of changing the ES can be modulated by controlling the cell-cell interaction (i.e., surface tension and cell adhesion), and vice versa. This finding provides a general concept that the amount and location of empty space can be a target for the regulation of cell arrangement, similar to the regulation of cell-cell interactions. This study proposes that in addition to the global feature of geometric constraints such as AR, local features such as the amount and location of the ES, play important roles in cell arrangement. These local features should consequently result in the regulation of cell functions.

## 2 Results

### 2.1 Development of a new cell morphology model to address unsolved questions

To clarify the questions in this study, we reviewed the current knowledge by re-investigating the images obtained in our previous research (Yamamoto and Kimura 2017). The 4-cell stage embryo of the wild-type *C. elegans* (aspect ratio AR≃1.7) shows the Diamond-type cell arrangement, which is robustly observed as the AR is decreased to as low as 1.2 by the *dpy-11* mutation (Fig. 1A(a1), 1B). When we increased the AR from the WT using the *lon-1* mutation and knockdown of the *C27D9*.*1* gene, we still observed the diamond-type arrangement in a robust manner, but the T-shaped and linear-types were also observed(Fig. 1A(a2),(a3) and 1B). We previously proposed the asymmetric attraction (AA) model that accounts for the robustness and diversity in the cell arrangement (Yamamoto and Kimura 2017), but this model had a limitation. The AA model predicted that the high AR embryos with impaired cell adhesion shows the linear type arrangement. However, when we knocked down the genes involved in cell adhesion (i.e., *hmr-1* and *hmp-2*), the embryos showed the T-shaped or T-reverse type arrangement (Fig. 1A(a4), 1B(right panel)). In contrast, the T-reverse type arrangement was never observed in the AA model.

Based on the discrepancy between the experimental observation and the consequences of the AA model, we set two questions to be addressed in this study. First, how can we explain the T-reverse type arrangement (question #1)? Second, why do embryos take different cell arrangements even when they have similar AR (question #2)? The coexistence of different types of cell arrangement within the same AR indicates that other aspects of geometrical constraints other than AR play important roles.

To find such new components of geometrical constraints, we chose an approach that could examine the effect of the precise geometry of the eggshell. We developed the cell morphology model using a multi-phase-field method (Fig. 1C) such that

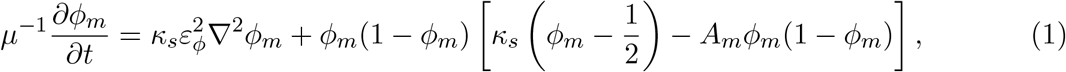

where

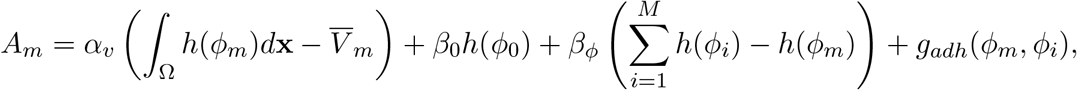

*ϕ*_0_ and *ϕ*_*m*_(*m* = 1, · · ·, 4) are the phase-field functions of the eggshell and cells, respectively. The first term of *A*_*m*_ defines the volumes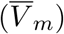 of *m*-th cell, the second and third terms of *A*_*m*_ define the repulsive condition between cell and eggshell and between cells, respectively. The fourth term, *g*_*adh*_(*ϕ*_*m*_, *ϕ*_*i*_), of *A*_*m*_ defines the attraction between *m*-th cell and *i*-th cell. *κ*_*s*_ is the parameter of surface tension (See Material and Methods for more details). We extended the model of Eq.(1) to a data-combined-model where the actual eggshell shapes from image data were directly combined with the eggshell phase-field function *ϕ*_0_ (Fig. 1D). The eggshell shapes used in the simulations are shown in Fig. 1E.

### 2.2 Reproduction of the T-reverse arrangement and sensitivity to eggshell geometry

We first asked #1: whether we can reproduce the T-reverse type arrangement by using the cell morphology model. The wild type *C. elegans* embryo shows a diamond arrangement at the 4-cell stage as shown in Fig. 2A (upper panels)(Movie S1). It is observed that after the division of AB and P1 cells, one of the cells (i.e., ABp cell) moves dynamically to adhere with the P2 cell and that the diamond arrangement settles down. In contrast, the T-reverse arrangement is found in the *hmr-1* ; *hmp-2* -double-knockdown strain with the *lon-1(e185)* mutant background where cell adhesion is impaired (Fig. 2B (upper panels), Movie S2). In the T-reverse arrangement, the ABp cell moves toward the posterior, whereas the movement of the EMS cell is smaller and does not adhere to the ABa cell.

**Figure 2:**
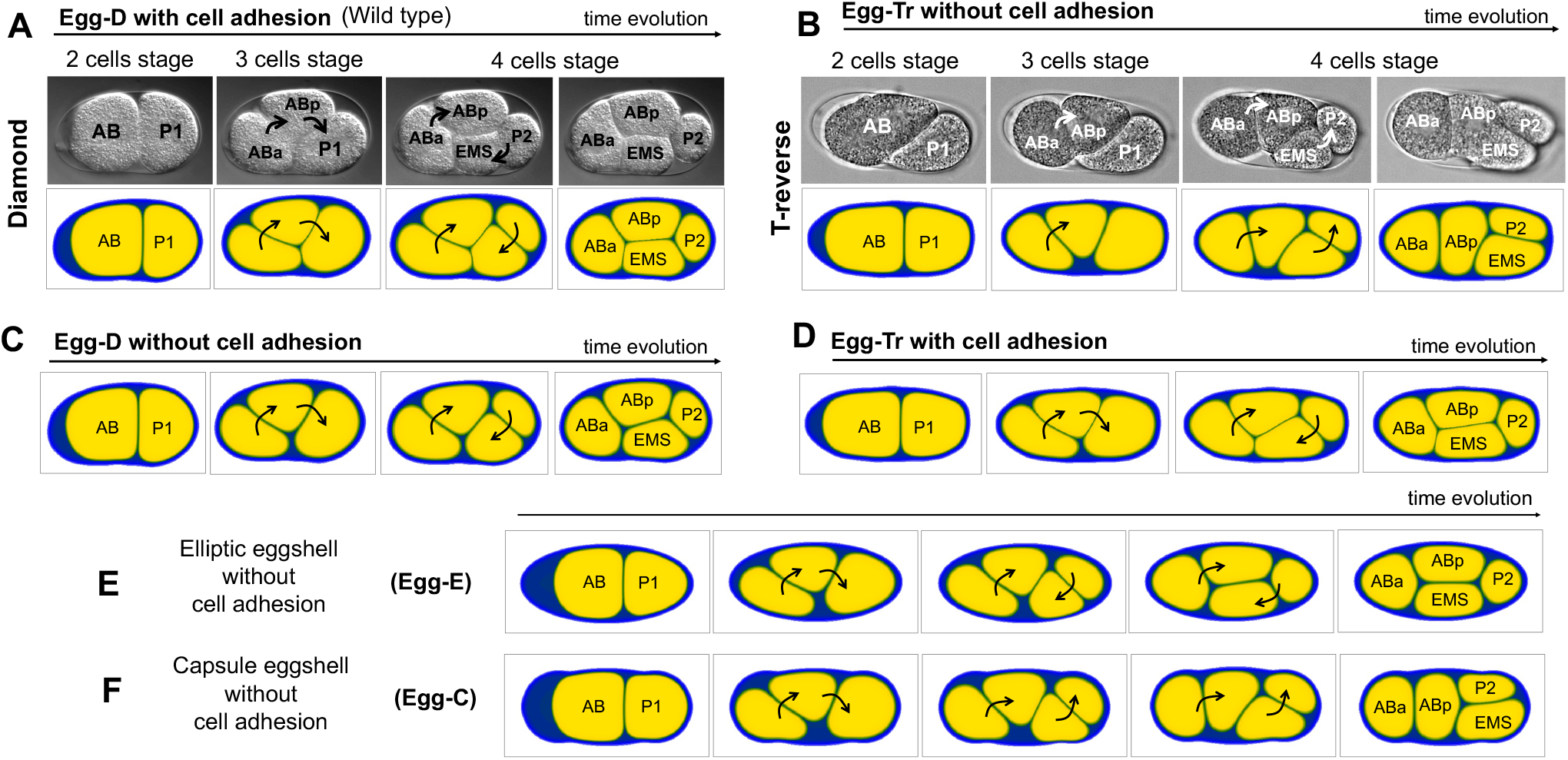
Geometrical effects for the T-reverse arrangement and representative simulation results. (A) Representative data for the diamond arrangement of the wild-type embryo (upper panels, Movie S1) and simulations (lower panels, Movie S3). Cell adhesions are present. The eggshell shape was generated from the image data of upper panels, and was the same as Egg-D(Fig. 1E). (B) Representative data for the T-reverse arrangement of the embryo (upper panels, Movie S2) and its simulations (lower panels, Movie S4). The embryo is that of a *hmr-1* ; *hmp-2* -double-knockdown strain with a *lon-1(e185)* mutant background, where cell adhesions were impaired. The eggshell shape was generated from the image data of upper panels, and was the same as Egg-Tr(Fig. 1E). (C) Simulation result for the case in which we removed the cell adhesions in (A). (D) Simulation result for the case in which we added cell adhesions in (B) (Movie S5). (E-F) Numerical experiments for the case in which the eggshells have the same AR (AR = 2.2) but different shapes. The eggshells, Egg-E in (E) and Egg-C in (F), were used(Fig. 1E). Black or white arrows indicate the direction in which the cells rotate.

We speculated that a characteristic of eggshell shape other than the AR is involved in inducing the T-reverse arrangement, as discussed by Yamamoto and Kimura (2017). First, to evaluate the contribution of the precise shape of the eggshell, we first regenerated actual eggshell shapes by converting the experimental image of the eggshell into the phase-field function *ϕ*_0_ and simulated the model of Eq.(1). We named the eggshell shape extracted from a wild-type embryo, which had an AR of 1.8, as Egg-D, whereas that from a *lon-1* mutant embryo with an AR of 2.1, was designated as Egg-Tr (Fig. 1E). We first tested the wild-type condition (i.e., Egg-D with cell adhesion), and successfully reproduced the diamond arrangement (Fig 2A(lower panels), Movie S3), supporting the feasibility of our model. Next, we incorporated the condition of the T-reverse arrangement (i.e., Egg-Tr without cell adhesion). Surprisingly, the T-reverse arrangement was successfully reproduced in our first trial (Fig 2B(lower panels), Movie S4). This was the first example of success in reproducing the T-reverse arrangement in a mathematical model. Further, the result suggested that the precision of eggshell geometry is critical for cell arrangement; this will be investigated further in a later section.

In the experiment, the T-reverse arrangement appeared only when cell adhesion was impaired (Yamamoto and Kimura 2017). To test the dependency on adhesion, we simulated a case with the shape of Egg-D without cell adhesion (Fig 2C), and of Egg-Tr with cell adhesion (Fig 2D, Movie S5). We found that the WT eggshell (Egg-D) took the diamond arrangement even without adhesion, whereas the T-reverse eggshell (Egg-Tr) with cell adhesion changed to the diamond arrangement. The consequences of the model were consistent with the experimental observations of *hmr-1;hmp-2* (RNAi) with normal eggshell, and of normal adhesion with an elongated eggshell (Yamamoto and Kimura 2017). This result supports the idea that cell adhesion (i.e., the attractive force between cells) plays an important role in improving the robustness of the diamond type of cell arrangement.

Why were we able to reproduce the T-reverse type arrangement, while our previous model in Yamamoto and Kimura (2017) could not? Most theoretical studies on *C. elegans embryos*, including Yamamoto and Kimura (2017), have assumed an ellipsoidal/elliptical shape of the eggshell. However, the actual shape of the eggshell, which we incorporated in our model, is more like a capsule, i.e., a tube with the two ends covered by two hemispheres. To test whether the difference between elliptical and capsule shapes was critical to reproduce the T-reverse type, we compared the elliptical (Egg-E) and capsule (Egg-C) shapes of eggshell with the same AR = 2.2 and without cell adhesion (Fig 2E, F). We found that the capsule shape induces the T-reverse arrangement, whereas the elliptical shape induces the diamond arrangement. This result supports the hypothesis discussed in Yamamoto and Kimura (2017) that the roundness of the eggshell edge might affect the generation of the T-reverse arrangement. The eggshell space in the posterior edge in the case of a capsule shape is wider than that of an elliptical shape, and therefore, EMS and P2 do not need to rotate much to align with the long axis of the eggshell, consequently leading to a T-reverse arrangement. In summary, the results demonstrate that certain components in the geometry of eggshell, other than the AR, are involved in determining cell arrangements.

### 2.3 The effect of extra-embryonic space on cell arrangement

What was the missing component of the geometrical constraints that enabled our present model to reproduce the T-reverse arrangement whereas the previous AA model could not? Such components might be included in the difference between the capsule and ellipsoidal shape. A hint to identify the specific factors involved in eggshell geometry is provided by the following analysis. We first focused on the parameter of cell surface tension (*κ*_*s*_) in our model of Eq.(1) and changed it to different scales. From the simulation tests, we found that the T-reverse arrangement can be obtained when the cell surface tension is increased even in the eggshell shape of the WT (Egg-D) (Fig. 3A). Furthermore, we found that the diamond arrangement can be obtained when the cell surface tension is decreased even in the eggshell shape of the T-reverse arrangement (Egg-Tr) (Fig. 3B). We presumed that the cell surface tension is likely to be effective when the cells are not compressed within the eggshell. These results suggested that the extra-embryonic space (ES) within the eggshell, i.e., the inner space of eggshell that is not occupied by the cells, may affect the cell arrangement.

**Figure 3:**
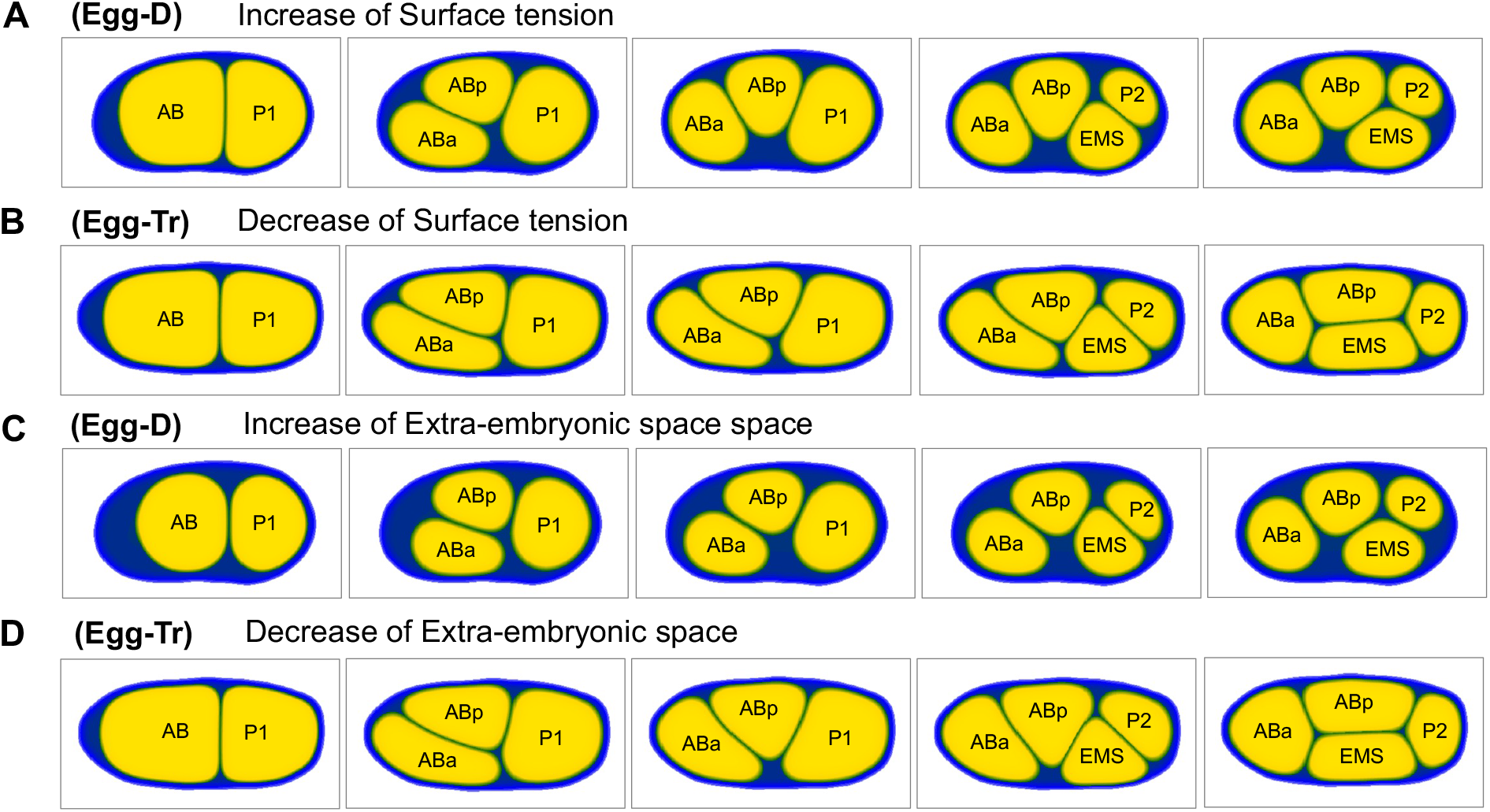
The effect of surface tension and empty space. Cell adhesions were not included in the (A-D) simulations. (A) The surface tension (*κ*_*s*_) was increased from the case of Fig. 2C. Egg-D (AR = 1.8) was used. (B) The surface tension (*κ*_*s*_) was decreased from the case of Fig. 2B. Egg-Tr (AR = 2.1) was used. (C) The extra-embryonic space was increased from the case of Fig. 2C. Egg-D (AR = 1.8) was used. (D) The extra-embryonic space was decreased from the case of Fig. 2B. Egg-Tr (AR = 2.1) was used.

We tested this idea by changing the ES in the model under the condition of fixed surface tension. We found the T-reverse arrangement when we increased the ES by decreasing the volume of cells but maintaining the relative size between the cells in the eggshell of WT (Fig. 3C, Egg-D). Similarly, we found a diamond arrangement when we decreased the empty space and compressed the cells in the eggshell of Egg-Tr (Fig. 3D, Egg-Tr). The results indicated that the ES of the eggshell plays as an important factor in determining cell arrangement.

### 2.4 Correlation between the extra-embryonic space and cell arrangement in experiments

In the previous section, we found that the extra-embryonic space (ES) can affect cell arrangement. To examine the feasibility of this idea, we analyzed the ratio of ES to the eggshell area for 230 *C. elegans* embryos with various ARs as obtained in our previous study (Yamamoto and Kimura 2017) (Fig. 4). Among embryos with similar AR, T-shaped and T-reversed type embryos tended to have a larger ES ratio (Fig. 4A). In other words, embryos with a large ES tended to take T-shape or T-reverse arrangements (Fig. 4B). These observations supported our idea that the amount of ES affects the cell arrangement even if the embryos have comparable ARs. Furthermore, the ES ratio and AR themselves had little correlation (Fig. 4C), indicating that ES can be considered as an independent factor of AR on geometrical constraints. In summary, based on the modeling and experimental supports, we concluded that the ES ratio can be an important geometrical constraint to determine cell arrangement in addition to the AR of the eggshell.

**Figure 4:**
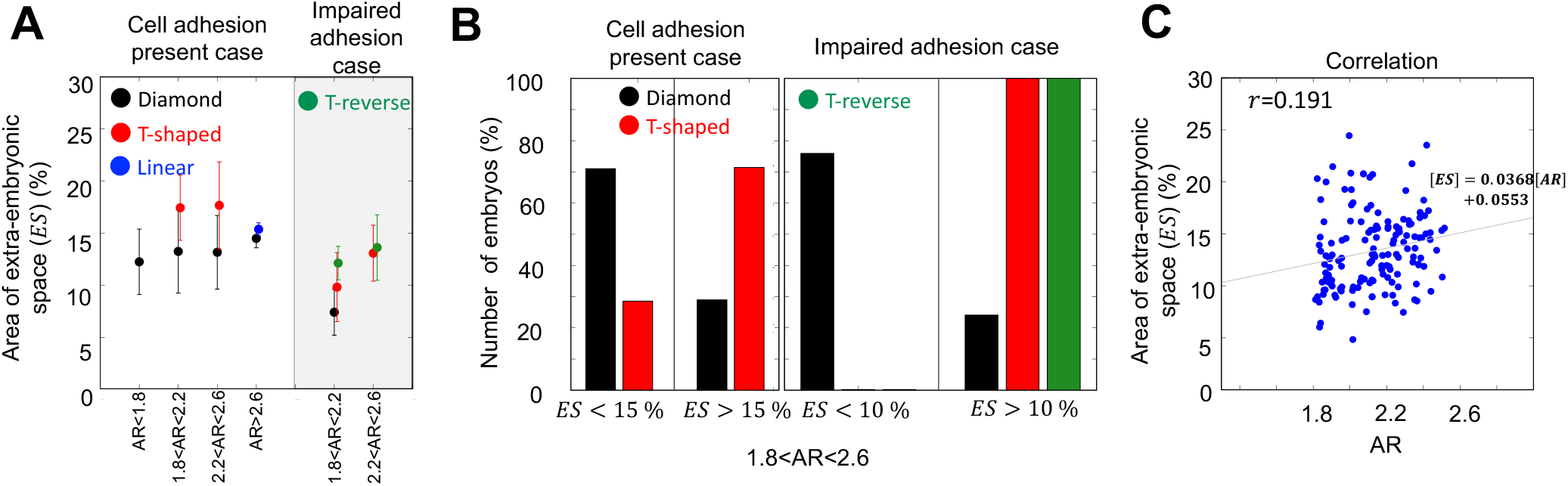
Analysis of extra-embryonic space in the *C. elegans* embryo. *N*_*s*_ = 230. (A) The ratio of extra-embryonic space (ES) for each type of cell arrangement in eggshells with similar AR. (B) The classification of embryo numbers by ES = 14%. Eggshells with 2.0 < *AR* < 2.6 were used. (C) The correlation between AR and ES. *r* is the correlation coefficient.

### 2.5 Diversity of cell arrangement induced by the ratio of the extra-embryonic space and eggshell shape

With regard to question #2 raised in the first section of the results, namely, why embryos can take different cell arrangements even when they have similar AR, our finding of the ES and experimental observations provided an answer. As the difference in cell arrangements correlates with the scale of the ES, we anticipated that the variety in the amount of ES can cause variety in cell arrangements with similar AR.

To investigate this possibility from a mathematical viewpoint, we changed the ratio of ES from 10% to 50% with several scales of the aspect ratio (Fig. 5A-C). Further, to determine the influence of sensitivity to eggshell shape, we also compared the elliptical and capsule eggshells (Egg-E and Egg-C) under the condition with the same AR both with and without cell adhesion.

**Figure 5:**
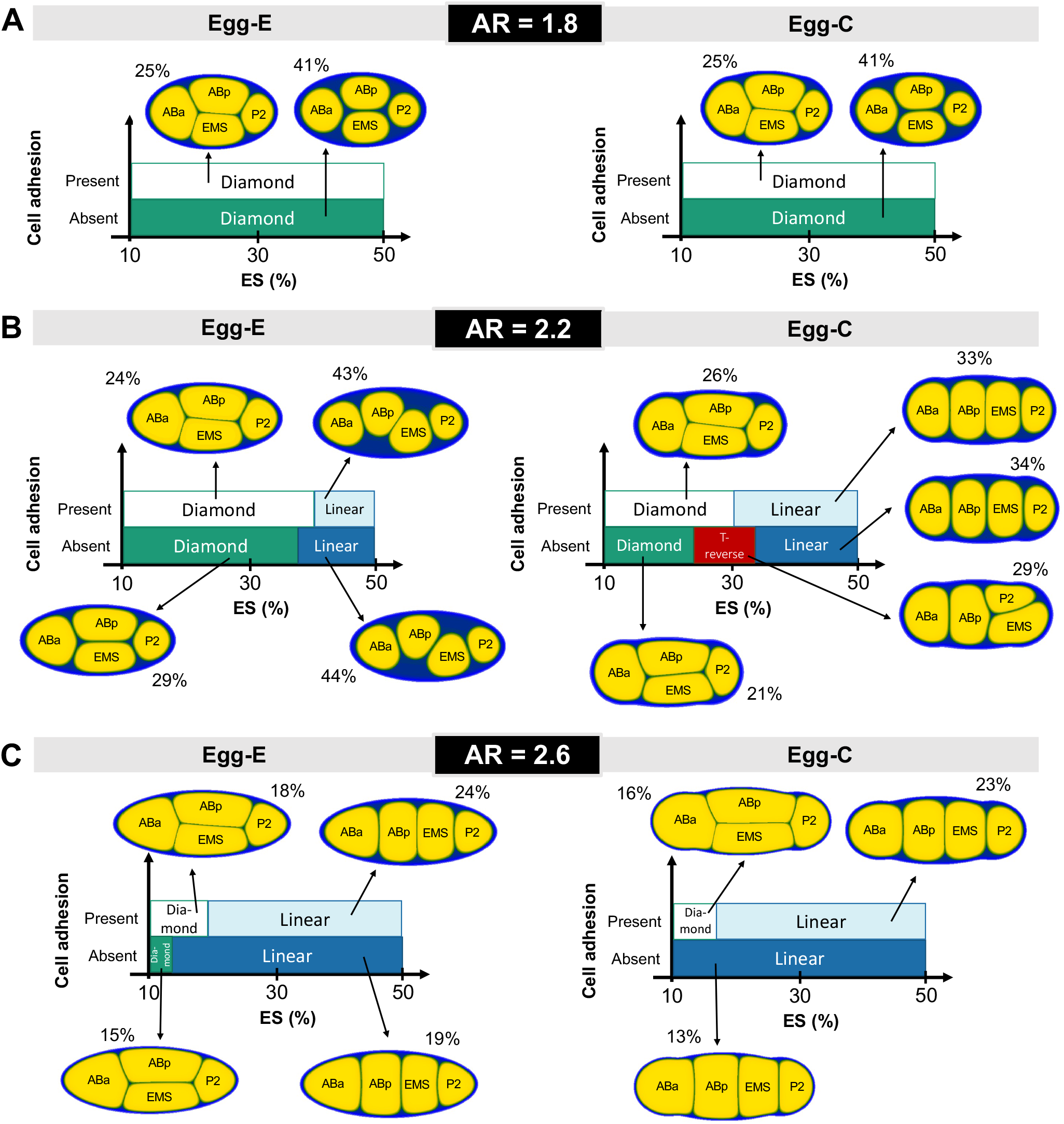
The effect of the extra-embryonic space ratio and eggshell geometry in cell arrangement. (A), (B), and (C) show the type of cell arrangement depending on the scale of the extra-embryonic space (ES) which was calculated by [1 − Total area of cells/Area of eggshell](%). The representative simulation results are shown in each case and [Number] % indicates the scale of the extra-embryonic space (ES).

First, the simulation results supported our idea that the amount of ES causes diversity in cell arrangement even with the same AR. We then focused on Egg-C with AR = 2.2 in the absence of cell adhesion (Fig. 5B, Egg-C, *Absent*), which is similar to the experimental condition of *lon-1; hmr-1; hmp-2* mutant/knockdown (Fig. 1A(a4), B(right)). The simulation predicted three types of arrangements (Diamond, T-reverse, and Linear) depending on the ES ratio. This result directly confirms that diverse arrangements can be reproduced within embryos having the same AR.

Further, we found that the sensitivity of cell arrangement against the ES ratio depends on the AR (Fig. 5B). When the AR is 2.2, as we explained in the previous paragraph, the cell arrangement was sensitive against the ES ratio (Fig. 5B). However, when the AR is 2.6, the sensitivity against the ES ratio was decreased. In both elliptical and capsule eggshells, the embryos showed the linear arrangement in a wide range of ES ratios and the ES ratio generating a diamond arrangement was very restricted (Fig. 5C). When the AR of the embryo is 1.8, the sensitivity to the ES is lost and the embryos take the diamond arrangement regardless of the ES ratio. Interestingly, the arrangement was also insensitive to the eggshell shape (i.e., Egg-E or Egg-C) and to the presence/absence of cell adhesion at AR = 1.8 (Fig. 5A).

Finally, we noticed that the effect of cell adhesion is also sensitive to ES. By comparing the presence or absence of cell adhesion, we found that cell adhesion can affect cell arrangement only in certain ranges of the ES ratio. For example, for Egg-C and AR = 2.2 (Fig. 5B, right), loss of cell adhesion changed the cell arrangement from the Diamond or Linear to the T-reverse when the ES was between ∼ 25 and ∼ 32%. In contrast, the cell arrangement remained unchanged regardless of cell adhesion for ES <∼ 25% and ES >∼ 32%.

Taken together, we concluded that the ES can be a critical factor to determine cell arrangement. The cell arrangement was sensitive to the amount of ES, and the effect of cell adhesion on cell arrangement was also sensitive to the ES. Meanwhile, the magnitude of sensitivity to cell arrangement with respect to the ES, the eggshell shape (i.e., elliptical or capsule), and cell adhesion, depended on the AR of the eggshell. This explains why different arrangements are found for the same AR in Fig. 1B. Therefore, a combinatorial effect of ES and AR underlies cell arrangement. Moreover, changes in the amount of ES might be a source of the variety in cell arrangement observed for various species of nematodes.

### 2.6 The effect of location of extra-embryonic space and T-shaped arrangement

In the previous section, we showed that the ES ratio can diversify cell arrangements even for the same AR. There remained two phenomena which we observed in the experiment (Fig. 4A, B) but did not reproduce in our model yet. One, we did not reproduce the T-shaped arrangement in our current simulations. Moreover, we could not explain why two different arrangements, T-reverse and T-shaped types, appeared even with a similar level of ES ratio and with similar AR. These limitations indicate the existence of additional factors that have not been considered in the model. We speculated that this missing factor is related to the property of the ES because the T-shaped arrangement was reproduced in the AA model (Yamamoto and Kimura 2017), and as the ES is the major difference between our multi-cellular morphology model and the AA model. Thus, we searched for the other property of ES, other than its amount, that determines whether the embryo takes the T-reverse or T-shaped arrangement.

Until now, in our model, we located the ES at the anterior side of the embryo as is often observed in experiments (Fig. 2A, B). We thus examined the effect of placing the ES in the opposite location, namely, the posterior side. Thus, we tested the effect of ES location in our model. In Fig. 6A, we placed the ES at the posterior side under the same condition as observed in the T-reversed arrangement shown in Fig. 2B (Egg-Tr eggshell without cell adhesion). We found that the embryo takes the T-shaped arrangement (the red square in Fig. 6A). The observed T-shaped arrangement was transient, and later, the simulated embryo took the diamond-type. Considering the rapid cell division occurring during the embryogenesis of *C. elegans*, the T-shaped arrangement observed *in vivo* might be the transient state at the time of cell division, before it reaches the steady state of cell arrangement. In a real embryo, the duration of the 4-cell stage is approximately 15 min (Yamamoto and Kimura 2017). In addition, our results indicated that both the T-reverse (Fig. 2B) and T-shaped (Fig. 6A) arrangements are reproduced with the common eggshell shape by changing the location of the ES. These results led us to propose that the diversity in location of the ES can explain the experimental result with the co-existence of the T-reverse and T-shaped arrangements in similar AR (Fig. 4B).

**Figure 6:**
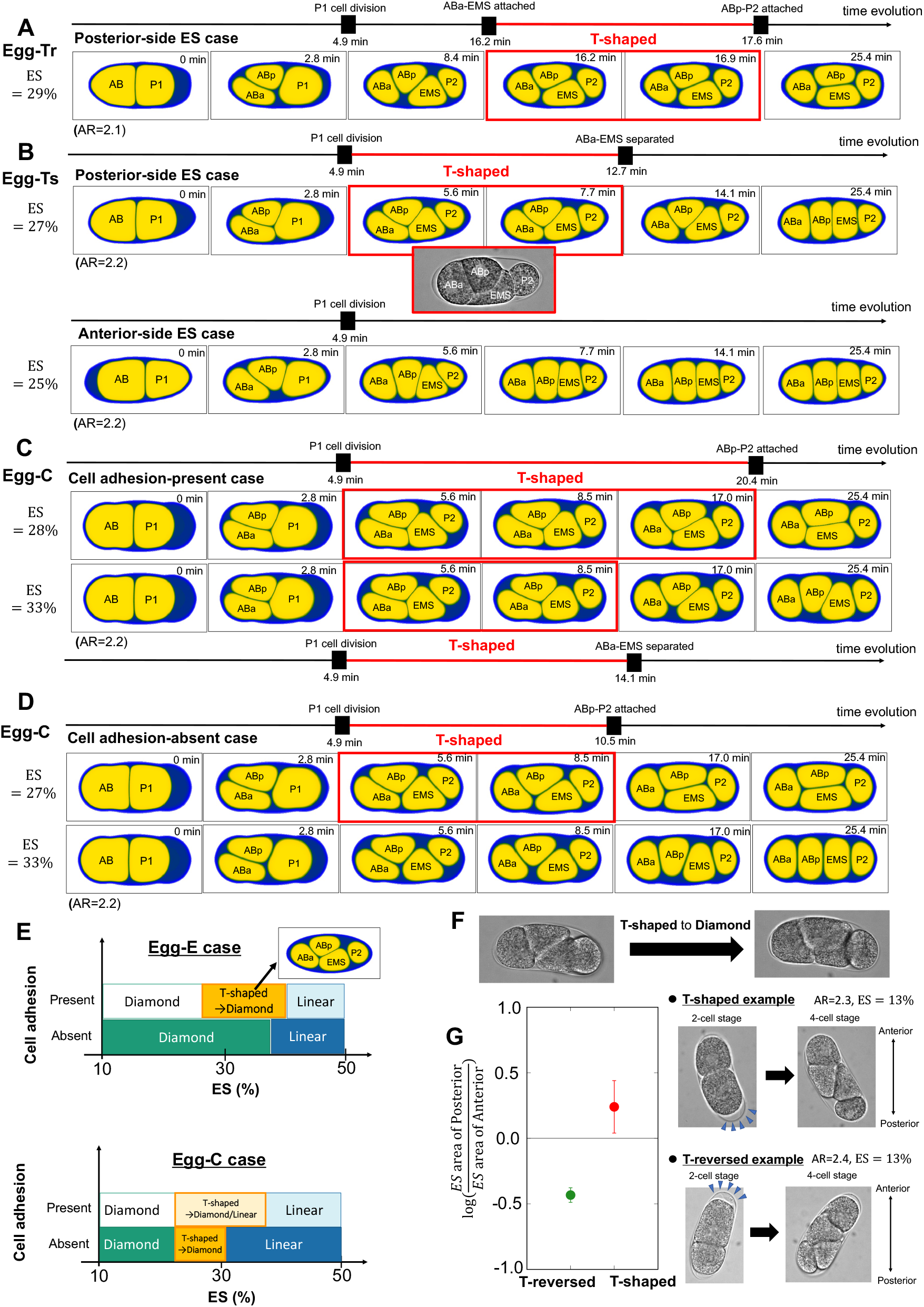
effect of the location of extra-embryonic space and the T-shaped arrangement. The location of extra-embryonic space (ES) is given in the posterior/anterior side of the eggshell in the two-cell stage. (A) Numerical experiments for posterior side ES under the T-reversed arrangement appearing in the experiment(Egg-Tr eggshell without cell adhesion) (B) Numerical experiments for the posterior and anterior sides of ES under the condition of T-shaped arrangement appearing in experiment(Egg-Ts eggshell with cell adhesion). The experimental figure of the T-shaped is the same as that in Fig. 1A(a2). (C-D) Representative simulations for cell adhesion present/absent cases on Egg-C eggshell shape. (E) The effect of extra-embryonic space amount in Egg-E and Egg-C eggshells with respect to posterior side ES. 1T4-shaped→Diamond/Linear indicates that the T-shaped arrangement is found in the intermediate phase and that it finally converges with the Diamond/Linear arrangement. (F) An example of an experiment where the T-shaped arrangement changed to the Diamond arrangement in an embryo with *lon-1; hmr-1; hmp-2* mutant/knockdown. (G) The data for [ES in the posterior]/[ES in the anterior] for T-reversed and T-shaped eggshells with *lon-1; hmr-1; hmp-2* mutant/knockdown. Right panels show the examples of each case.

We next examined whether we could also reproduce the T-shaped arrangement in the presence of cell adhesion. In the experiments, we frequently observed the T-shaped arrangement in the presence of cell adhesion (Fig. 1A(a2)). Therefore, we used the condition of the Egg-Ts eggshell with cell adhesion, and placed the ES at the posterior/anterior side (Fig. 6B). We succeeded in reproducing the T-shaped arrangement when we placed the ES in the posterior side, whereas the final arrangement became the linear type (Fig. 6B, upper panels). In contrast, the T-shaped arrangement was not observed at any time point when the ES was placed in the anterior side (Fig. 6B, lower panels). These results indicate that the location of the ES is critical to reproduce the T-shaped arrangement and is thus, an additional factor that affects cell arrangement.

Next, we systematically examined how the cell arrangement is affected by the amount of the ES in the presence and absence of cell adhesion and in Egg-C and Egg-E shapes when the ES is located at the posterior side (Fig. 6C-E), as we investigated in Fig. 5B for the ES location on the anterior side. We found that the T-shaped arrangement appears in the intermediate phase and that the final arrangement is either the Diamond or Linear type (Fig. 6C,D), as in the results of Fig. 6A,B. We also found that an increase in the ES ratio decreases the duration of the T-shaped state (Fig. 6C). Furthermore, we found that the state of the T-shaped arrangement is less likely to be generated in the absence of adhesion compared to in its presence (Fig. 6C and D). Indeed, we found several experimental examples showing the appearance of the T-shaped arrangement, but it finally converges to the Diamond arrangement in embryos with impaired cell adhesion (Fig. 6F).

Fig. 6E shows that the ES ratio consistently affects the cell arrangement, so that the T-shaped arrangement is not formed either when the size of the ES is small or when it is large. Meanwhile, we did not find the T-shaped arrangement for any size of ES on the Egg-E eggshell when cell adhesion was absent. This finding is consistent with the experimental observation that the T-shaped arrangement is observed more frequently when cell adhesion is present (Yamamoto and Kimura 2017). This result also suggests that the precise shape of the eggshell (e.g., the difference between Egg-E and Egg-C) is critical to generate the T-shaped arrangement when cell adhesion is absent.

Finally, we examined the experimental data of whether the ES is located at the anterior/posterior side at the 2-cell stage embryos with *lon-1; hmr-1; hmp-2* mutant/knockdown and AR = 2.3 or 2.4 which result in T-reversed or T-shaped arrangements at the 4-cell stage (Fig. 6G). We plotted the data of log([ES area in posterior edge]/[ES area in anterior edge]) for 6 embryos. The results showed that the T-reversed case has more ES in the anterior side, whereas the T-shaped case has more ES in the posterior side. This experimental observation is consistent with our idea that the location of ES is critical for the choice between the T-reversed and T-shaped arrangements under conditions of impaired cell adhesion with similar AR.

## 3 Discussion

### 3.1 Construction of cell morphology model by using multi-phase-field method

Application of the multi-phase-field model for multi-cellular systems has been introduced by Nonomura (2012), and this modeling method has been applied in various biological phenomena (Akiyama et al. 2018; Moure and Gomez 2021; Seirin-Lee et al. 2020; Seirin-Lee 2016). In this study, we aimed to investigate the role of spatial constraints in the cell arrangement of *C. elegans* embryos from a geometrical viewpoint. For this purpose, we adopted a phase-field method. We traced the actual eggshell shape of the *C. elegans* embryo, and incorporated it into the phase-field model. The phase-field model is advantageous for describing the precise shape of the eggshell; thus, we did not need to approximate the shape by using a simplified shape such as an ellipsoid.

In this study, we adopted a 2-dimensional model rather than a 3-dimensional model, to intensively incorporate the precise shape of the eggshell traced from the experimental data and to find unknown geometrical effects by examining various parameter values with reduced numerical costs. However, mathematical formulation of phase-field modeling is written in general as an *N*-dimensional spatial format, so that extension to 3-dimensional is simply possible once the obstacle of numerical costs is solved. Notably, the 2-dimensional model is sufficient to examine the diamond, T-shaped, T-reverse, and linear types of cell arrangement at the 4-cell stage, because the centers of the 4 cells are roughly aligned on a common plane. For other cases, such as the pyramid-type arrangement or arrangement in later stages, 3-dimensional modeling will be required. In future, systematic analysis using 3-dimensional imaging and modeling is expected to contribute to a detailed understanding regarding the role of spatial constraints.

### 3.2 Reproduction of the T-reverse type cell arrangement

The specific motivation of this study was that we were not able to reproduce the T-reverse type cell arrangement in our previous research (Yamamoto and Kimura 2017). Using our cell morphology model, we succeeded in reproducing the T-reverse type arrangement (Fig. 2B). Incorporation of the actual eggshell shape was a key factor (Fig. 2). Further, we narrowed down that the amount (Fig. 5) and location (Fig. 6) of the extra-embryonic Space (ES) was critical to reproduce the T-reverse arrangement.

### 3.3 The amount and location of the extra-embryonic space (ES) as the cause of diversity in cell arrangement

Through this study, we revealed that the cell arrangement varies depending on the amount and location of the ES even under a fixed cell division orientation, cell-cell interaction, and the AR of the geometric constraint. This raises an important message that we should consider the state of the ES (or equivalent empty spaces) when we consider the mechanisms underlying the cell arrangements.

We found that the change in the amount of the ES had a similar effect as the change in cell surface tension (Fig. 3). Interestingly, the extracellular environment and cell autonomous activity plays an interchangeable role in cell arrangement. The nematode species show diverse cell arrangement even if they have a similar aspect ratio of the eggshell (Yamamoto and Kimura 2017). Previously, we speculated that this difference is caused by the difference in cell adhesion/tension(Yamamoto and Kimura 2017). The present study provides another possibility that the change in the amount or location of the ES causes diversity among species.

In Fig. 7, we examined the amount of ES in other nematode species. We investigated the cell arrangement types in 5 families from the data of Dolinski et al. (2001) (Fig. 7A), indicating that different families have different cell arrangement types that appear predominantly. *Rhabditina*(①), *Diplogastrina*(②), and *Panagrolaimidae*(③) predominantly show a diamond arrangement. In contrast, *Cephalobidae*(④) and *Tylenchina*(⑤) predominantly show T-shaped and Linear arrangements, respectively. Interestingly, when we classified these different families by the ES ratio, we found that *Rhabditina*(①), *Diplogastrina*(②), and *Panagrolaimidae*(③) mainly show ES< 18% whereas *Cephalobidae*(④) and *Tylenchina*(⑤) mainly show ES> 18% (Fig. 7B). This observation indicates that a higher ratio of ES is observed in the case of T-shaped or Linear arrangements. Indeed, we found that the arrangement types can be classified by the same measure of ES ratio (Fig. 7C), proposing that ES ratio may be an important measure to induce the different cell arrangements for the different families of nematodes.

**Figure 7:**
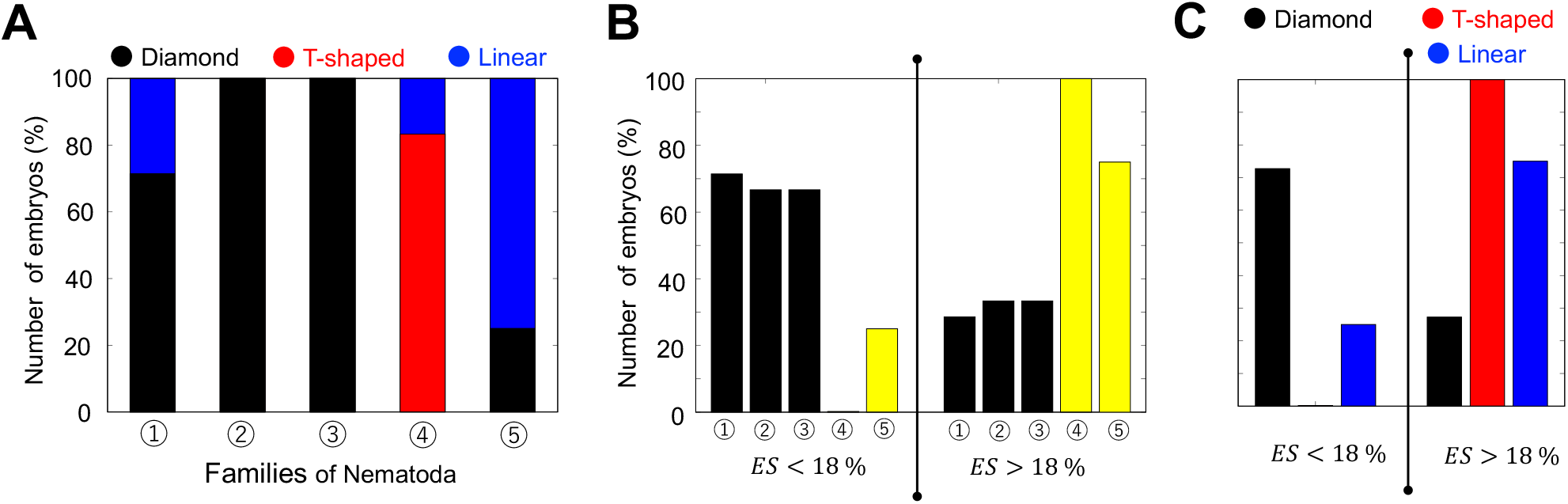
Analysis of cell arrangement types and extra-embryonic space in several families of nematodes. Image data of eggshells with 1.5 < *AR* < 2.7 in Dolinski et al. (2001) was used for the analysis. ①:*Rhabditina* (*N*_*s*_ = 7), ②:*Diplogastrina* (*N*_*s*_ = 3), ③:*Panagrolaimidae* (*N*_*s*_ = 3), ④:*Cephalobidae* (*N*_*s*_ = 6), ⑤:*Tylenchina* (*N*_*s*_ = 4). (A) The ratio of cell arrangement types for several families of nematodes; Diamond(black), T-shaped(red), and Linear(blue). (B) Classification of embryo numbers for each family of nematodes by ES = 18%. (C) Classification of embryo numbers for each cell arrangement type by ES = 18%.

However, although the amount of the ES is important, it is not the sole determinant of cell arrangement. This is because the data in Fig. 7 also includes exceptional cases. The diversity among species might be caused by a combination of other factors such as cell division orientation, cell-cell interactions, and the AR of the geometric constraints.

While the ES and cell adhesion/tension introduce diversity in cell arrangement, we found that some aspect ratios, such as AR = 1.8, result in a specific cell arrangement regardless of the amount of the ES or the presence of cell adhesion (Fig. 5A). Interestingly, this AR matches the AR of the wild-type *C. elegans*. Moreover, all nematode species with AR = 1.8 investigated so far only take the diamond arrangement at the 4-cell stage(Yamamoto and Kimura 2017). Thus, AR = 1.8 may be suitable for the robust arrangement, and might thus be adopted in nature.

### 3.4 The role of geometric constraints in the determination of cell arrangements

Cell arrangement is important for development and is known to be determined by three factors: the orientation of cell division, strength of cell-cell interaction (i.e., attraction and repulsion between the adjacent cells), and geometric constraints. Compared to the first two factors, investigations on geometric constraints have been limited. Previously, we demonstrated the contribution of the aspect ratio (AR), a global feature of geometric constraints (Yamamoto and Kimura 2017). In this study, we demonstrated the contribution of empty space, which is a local feature of geometric constraints. The empty space is the extra-embryonic space (ES) in the *C. elegans* embryo. We believe that the amount and location of empty space is also important for cell arrangement in other cell types of cell stages later than the 4-cell stage, and in species other than *C. elegans*. This research is currently underway.

## 4 Materials and Methods

To describe the shapes of cells precisely, we chose phase-field modeling. The technical application of the phase-field method for the morphodynamics of single cells is well-introduced in Shao et al. (2010, 2012). On the contrary, the multi-phase field method applied for the multi-cellular system is well-described in Nonomura (2012), and its applications to pattern formation in the multicellular system and nuclear chromatin dynamics are shown in Seirin-Lee (2016); Seirin-Lee et al. (2016, 2020). In this paper, we constructed a mathematical model describing the process from the 2-cell stage to 4-cell stage of the *C. elegans* embryo by using the multi-phase field method introduced by Nonomura (2012), Seirin-Lee et al. (2016), and Akiyama et al. (2018). Note that the phase-field model description is basically written in the *N*-dimensional spatial case, and there is no difference in the formulation of modeling between high-dimensional cases.

### 4.1 Multicellular morphology model of *C. elegans* embryo

We designed the model by using phase-field functions defined by the eggshell, *ϕ*_0_(**x**), and the daughter cells derived from the fertilized mother cell, *ϕ*_*m*_(**x**, *t*) ∈ [0, 1] (1 ≤ *m* ≤ *M*), where **x** ∈ Ω in **R**^*N*^, *t* > 0, *M* is the total number of cells, and Ω is the area of the system. In the 2-cell stage, *M* = 2 and the AB and P1 cells are defined by the first cell (*m* = 1) and second cell (*m* = 2). In the 4-cell stage, *M* = 4 and ABa and EMS cells are defined by the first cell (*m* = 1) and second cell (*m* = 2), ABp and P2 cells are defined by the third cell (*m* = 3) and fourth cell (*m* = 4), respectively (Fig. 1C). The regions of the cells and eggshell are defined by {**x**|0 *< ϕ*_*m*_(**x**, *t*) ≤ 1} and {**x**|0 < 1 − *ϕ*_0_(**x**) ≤ 1}, respectively.

The detailed model description by using energy functionals is described in the Supplementary information. Here, we briefly describe the transformed evolutionary system that follows. Through transformation to the evolutionary system from the energy functionals (Supplementary information), the model equation of the eggshell and the time evolution of the *m*-th cell is given by the following equations:

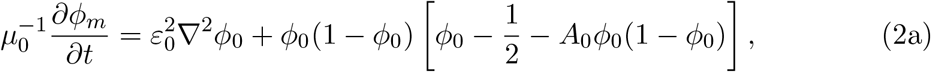

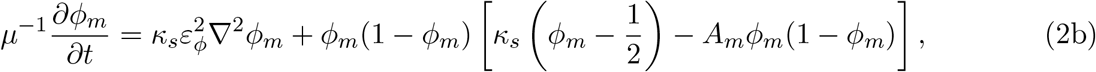

where *µ*_0_, *µ, ε*_0_, *ε*_*ϕ*_ are positive constants and *κ*_*s*_ is the surface tension. *A*_0_ and *A*_*m*_ are given by

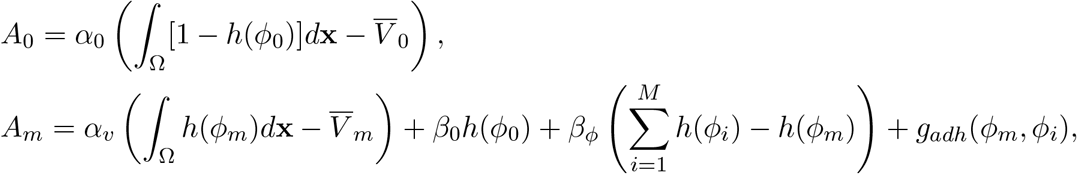

where *α*_0_, *α*_*v*_, *β*_0_, *β*_*ϕ*_ are positive constants. 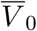 is the volume of the eggshell, 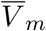 is the volume of *m*-th cell, and *h*(*ϕ*) = *ϕ*^3^(10 − 15*ϕ* + 6*ϕ*^2^). The first terms of *A*_0_ and *A*_*m*_ define the volumes of the eggshell and *m*-th cell. The second and third terms of *A*_*m*_ define the territory (repulsive) condition in which the cell regions are not overlapped. The fourth term, *g*_*adh*_(*ϕ*_*m*_, *ϕ*_*i*_), of *A*_*m*_ defines the attraction between *m*-the cell and *i*-th cell and is given as

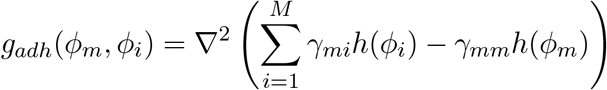

where *γ*_*mi*_ is the strength of attraction between the *m*-th cell and *i*-th cell so that *γ*_*mi*_ = *γ*_*im*_. As the ABa and P2 cells are not in contact in the 4-cell stage, we set *γ*_14_ = *γ*_41_ = 0. Based on the experiment of Yamamoto and Kimura (2017), we assumed that the attractions, EMS– P2 (*γ*_24_) < ABa–ABp (*γ*_13_) < ABa–EMS (*γ*_12_) ≈ ABp–EMS (*γ*_23_). As there are no data for the attraction between ABp cells and P2 cells (*γ*_34_), we used a similar scale for the attraction between the EMS cell and P2 cell (*γ*_24_).

### 4.2 Cell division in sequence

The direction and position of the cell division plane is an important element that can affect the initial cell position. Spindle positioning, which determines the location of the cell division plane, is tightly regulated by both biochemical and biophysical dynamics (Coffman et al. 2016; Cheng et al. 1995; Kimura and Onami 2007; Minc et al. 2011; Manuel Théry et al. 2007). However, because the focus of this study is on how cell arrangement is affected by the geometrical effects of the eggshell, we fixed the direction and position of the cell division plane based on the image of the wild-type *C. elegans* embryo.

We thus considered the simplest model where the direction and position of the cell division plan is determined by the plane bisecting the one connecting the given spindle poles at the center (Fig. 1C) (Akiyama et al. 2018). Let us set the spindle poles of *m*-th cell by 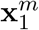 and 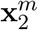 and the divided daughter cells of *m*-th cell, *ϕ*_*m*_, by *ϕ*_*m*,1_, and *ϕ*_*m*,2_. Then the daughter cells are described as:

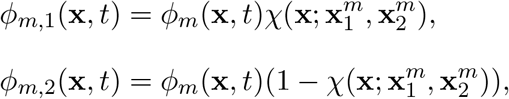

where

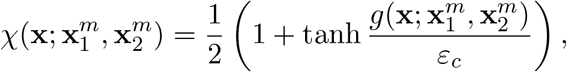

and *ε*_*c*_ is a positive constant. The function of *χ* is the step function which has 1 or 0 depending on the region bisecting the division plan, 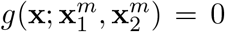. The function of 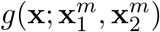 is defined as:

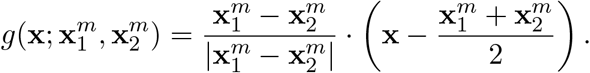

We started simulations with the 2-cell stage of the AB cell (*ϕ*_1_) and P1 cell (*ϕ*_2_). When *t* = *t*_1_, we first divided the AB cell to the ABp cell (*ϕ*_1,1_) and ABa cell (*ϕ*_1,2_). Then we divide P1 cell to P2 cell (*ϕ*_2,1_) and EMS cell (*ϕ*_2,2_) when *t* = *t*_2_(> *t*_1_). After cell division, we replaced the phase-field functions with *ϕ*_1,1_(**x**, *t*) = *ϕ*_3_(**x**, *t*), *ϕ*_1,2_(**x**, *t*) = *ϕ*_1_(**x**, *t*), *ϕ*_2,1_(**x**, *t*) = *ϕ*_4_(**x**, *t*), and *ϕ*_2,2_(**x**, *t*) = *ϕ*_2_(**x**, *t*).

### 4.3 Regeneration of the actual eggshell shape and parameter values

To investigate the geometric effect of the eggshell precisely, we simulated a model (2b) with *ϕ*_0_(**x**) that reflects the actual shape of the *C. elegans* eggshell by converting the images from the experiment to numeric data (Fig. 1D). Conversion of the eggshell image to numeric data and the imaging analysis were performed using ImageJ (Version:2.1.0/1.53c).

The eggshell volume was fixed for all the simulations, and the volume of each cell was determined based on the experimental data (Table S1). *ε*_0_ and *ε*_*ϕ*_ were estimated from the thickness of the eggshell (*δ*_0_ = 0.4 *µm*) (Krenger et al. 2020) and cell membrane (*δ*_*ϕ*_ = 0.01 *µm*) (Alberts et al. 2014) by using the formula for the interface width of the phase-field function (Seirin-Lee et al. 2016) such that

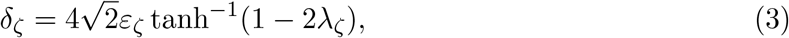

where *ζ* = {0, *ϕ*}, and *λ*_*ζ*_ is the value defining the interface region of the eggshell by *λ*_*ζ*_ *< ϕ*_*ζ*_(**x**) < 1 − *λ*_*ζ*_. The detailed values of *ε*_0_ and *ε*_*ϕ*_ are given to *ε*_0_ = 3.07764 × 10^−2^ and *ε*_*ϕ*_ = 4.172716 × 10^−3^ by choosing *λ*_0_ = 0.01 and *λ*_*ϕ*_ = 0.3 which defines the regions of the cells as *ϕ*_*m*_(**x**, *t*) > *λ*_*ϕ*_.

The spatial size and temporal scale were estimated by comparing the quantitative and qualitative dynamics of the cell arrangement in simulations with live imaging data from the 2-cell and 4-cell stages of *C. elegans* embryos (Fig. S1A and Table S1) From the experimental data, we approximated the interval of the cell division time to 4 minutes. Using these data, we estimated the time scale of the model as *t* = 14 seconds. The area of system was Ω = 95.5 *µm*×47.25 *µm* as estimated by the wild type *C. elegans* embryo size (long diameter= 54 *µm* and short diameter= 29.6 *µm*) and the non-dimensional numerical space Ω = [0, 1.8] × [0, 0.9]. The cell volumes (*V*_*m*_) were estimated as a relative scale compared to eggshell size, based on experimental data (Fig. S1B and Table S1).

The other parameter values for the phase-field functions were chosen by *µ*_0_ = 0.135, *µ* = 1.0, *α*_0_ = 200, *α*_*v*_ = 300, *β*_0_ = 50, *β*_*ϕ*_ = 30. The strength of the attraction between the cells were chosen to *γ*_24_ = *γ*_34_ = 0.002, *γ*_13_ = 0.003, *γ*_12_ = 0.008, *γ*_23_ = 0.006. The surface tension, *κ*_*s*_, was varied in [0, 13]. Note that the scales of these kinetic parameters in the phase-field functions are determined relatively depending on the interface values of the phase-field function. Since we have estimated the parameters affecting the interface dynamics of the phase-field function (the thickness of eggshell and cell membrane) from experimental data, our simulations can represent the results within the biologically feasible range of parameter values.

### 4.4 Microscopy observation of *Caenorhabditis elegans* embryos

In this study, we used the microscopy images of the *C. elegans* embryos obtained in our previous study Yamamoto and Kimura (2017). The methods used to obtain these images are described in the paper. Briefly, phase contrast images were acquired at room temperature under an inverted microscope (Axiovert 100; Carl Zeiss, Oberkochen, Germany) equipped with a 40 ×, 0.70 N.A. objective (Plan-Neofluar; Carl Zeiss) and a CCD camera (ORCA-100; Hamamatsu, Japan). To change the aspect ratio of the eggshell, the mutants of *dpy-11* and *lon-1* genes were used in combination with the RNAi-mediated gene knockdown of *C27D9*.*1* gene. The cell arrangement of the T-reverse type were observed when we knocked down the *hmr-1* and *hmp-2* genes by using RNAi, which encodes the proteins responsible for cell adhesion.

## Acknowledgements

We thank Kazunori Yamamoto (present address: Kanagawa Institute of Technology) for providing image data and for discussion. This work was supported by a Grant-in-Aid for Scientific Research from the Ministry of Education, Culture, Sports, Science and Technology, Japan, to S.S.L. (JP19H01805 and JP17KK0094) and to A.K. (JP19H01805 and JP18H02414).

## References

Akiyama, M., Nonomura, M., Tero, A., Kobayashi, R., 2018. Numerical study on spindle positioning using phase field method. Physical Biology 16, 016005.

Akiyama, M., Tero, A., Kobayashi, R., 2010. A mathematical model of cleavage. Journal of Theoretical Biology 264 (1), 84–94.

Alberts, B., Johnson, A., Lewis, J., Raff, M., Roberts, K., Walter, P., 2014. Molecular Biology of The Cell. Garland Science, New York.

Baena-López, L. A., Baonza, A., Garćia-Bellido, A., 2005. The orientation of cell divisions determines the shape of Drosophila organs. Current Biology 15, 1640–1644.

Bowerman, B., Tax, F. E., Thomas, J. H., Priess, J. R., 1992. Cell interactions involved in development of the bilaterally symmetrical intestinal valve cells during embryogenesis in Caenorhabditis elegans. Development (Cambridge, England) 116 (4), 1113–22.

Cheng, N. N., Kirby, C. M., Kemphues, K. J., 1995. Control of cleavage spindle orientation in Caenorhubditis elegans: The role of the genes par-2 and par-3. Genetics 139, 549–559.

Coffman, V. C., A, M. B. McDermottb Shtyllac, B., Dawes, A. T., 2016. Stronger net posterior cortical forces and asymmetric microtubule arrays produce simultaneous centration and rotation of the pronuclear complex in the early Caenorhabditis elegans embryo. Molecular Biology of the Cell 27 (7), 3550–3562.

Dolinski, C., Baldwin, J. G., Thomas, W. K., 2001. Comparative survey of early embryogenesis of Secernentea (nematoda), with phylogenetic implications. Canadian Journal of Zoology 79, 82–94.

Fickentscher, R., Struntz, P., Weiss, M., 2013. Mechanical cues in the early embryogenesis of Caenorhabditis elegans. Biophysical Journal 105, 1805–1811.

Gilbert, S. F., Michael, J. F., 2019. Developmental Biology, 12th edn. Sinauer Associates Inc., Sutherland, MA.

Gloerich, M., Bianchini, J. M., Siemers, K. A., Cohen, D. J., Nelson, W. J., 2016. Cell division orientation is coupled to cell-cell adhesion by the E-cadherin/LGN complex. Nature Communications 8, 13996.

Gönczy, P., 2005. Asymmetric cell division and axis formation in the embryo. WormBook.org, doi/10.1895/wormbook.1.30.1.

Kajita, A., Yamamura, M., 2002. Physical modeling of the cellular arrangement in C. elegans early embryo: Effect of rounding and stiffening of the cells. Genome Informatics 13, 224–232.

Kimura, A., Onami, S., 2007. Local cortical pulling-force repression switches centrosomal centration and posterior displacement in C. elegans. Journal of Cell Biology 179 (7), 1347–1354.

Krenger, R., Burri, J. T., Lehnert, T., Nelson, B. J., Gijs, M. A. M., 2020. Force microscopy of the Caenorhabditis elegans embryonic eggshell. Microsystems & Nanoengineering 6 (29), 0–11.

Manuel Théry, A. J.-D., Racine, V., Bornens, M., J’ulicher, F., 2007. Experimental and theoretical study of mitotic spindle orientation. Nature 447 (24), 493–497.

Mickey, K. M., Mello, C. C., Montgomery, M. K., Fire, A., Priess, J. R., 1996. An inductive interaction in 4-cell stage C. elegans embryos involves APX-1 expression in the signalling cell. Development (Cambridge, England) 122 (6), 1791–8.

Minc, N., Burgess, D., Chang, F., 2011. Influence of cell geometry on division-plane positioning. Cell 144, 414–426.

Moure, A., Gomez, H., 2021. Phase-field modeling of individual and collective cell migration. Archives of Computational Methods in Engineering 28, 311–344.

Nonomura, M., 2012. Study on multicellular systems using a phase field model. PLoS One 7, e33501.

Pierre, A., Sallë, J., Wühr, M., Minc, N., 2016. Generic theoretical models to predict division patterns of cleaving embryos. Developmental Cell 39, 667–682.

Schulze, J., Schierenberg, E., 2011. Evolution of embryonic development in nematodes. EvoDevo 2 (18), 1–16.

Seirin-Lee, S., 2016. Lateral inhibition-induced pattern formation controlled by the size and geometry of the cell. Journal of Theoretical Biology 404, 51–65.

Seirin-Lee, S., Osakada, F., Takeda, J., Tashiro, S., Kobayashi, R., Yamamoto, T., Ochiai, H., 2020. Role of dynamic nuclear deformation on genomic architecture reorganization. PLOS Computational Biology 15 (8), e1007289.

Seirin-Lee, S., Tashiro, S., Awazu, A., Kobayashi, R., 2016. A new application of the phase-field method for understanding the reorganization mechanisms of nuclear architecture. Journal of Mathematical Biology 74, 333–354.

Shahbazi, M. N., 2020. Mechanisms of human embryo development: from cell fate to tissue shape and back. Developmental 147, dev190629.

Shao, D., Levine, H., Rappel, W.-J., 2012. Coupling actin flow, adhesion, and morphology in a computational cell motility model. PNAS 109 (18), 6851–6856.

Shao, D., Rappel, W. J., Levine, H., 2010. Computational model for cell morphodynamics. Physical Review Letter 105, 108104.

Yamamoto, K., Kimura, A., 2017. An asymmetric attraction model for the diversity and robustness of cell arrangement in nematodes. Development 144, 4437–4449.

